# Improving genomically recoded *Escherichia coli* for the production of proteins containing non-canonical amino acids

**DOI:** 10.1101/2021.12.10.472167

**Authors:** Jessica G. Perez, Erik D. Carlson, Oliver Weisser, Camila Kofman, Kosuke Seki, Benjamin J. Des Soye, Ashty S. Karim, Michael C. Jewett

## Abstract

A genomically recoded *Escherichia coli* strain that lacks all amber codons and release factor 1 (*C321.ΔA*) enables efficient genetic encoding of chemically diverse, non-canonical amino acids (ncAAs) into proteins. While *C321.ΔA* has opened new opportunities in chemical and synthetic biology, this strain has not been optimized for protein production, limiting its utility in widespread industrial and academic applications. To address this limitation, we describe the construction of a series of genomically recoded organisms that are optimized for cellular protein production. We demonstrate that the functional deactivation of nucleases (e.g., *rne, endA*) and proteases (e.g., *lon*) increases production of wild-type superfolder green fluorescent protein (sfGFP) and sfGFP containing two ncAAs up to ∼5-fold. Additionally, we introduce a genomic IPTG-inducible T7 RNA polymerase (T7RNAP) cassette into these strains. Using an optimized platform, we demonstrated the ability to introduce 2 identical N_6_-(propargyloxycarbonyl)-_L_-Lysine residues site specifically into sfGFP with a 17-fold improvement in production relative to the parent. We envision that our library of organisms will provide the community with multiple options for increased expression of proteins with new and diverse chemistries.

## Introduction

The genetic code is a universal cipher that describes how mRNA codons are translated into proteins. Of the 64 available codons, 61 encode the twenty standard amino acids with the remaining three (UAA, UAG, UGA) responsible for signaling termination of protein synthesis.^[1]^ This biochemical principle extends through all kingdoms of life and for a long time was considered immutable. However, by the late 1970s variations in the genetic code, such as the reassignment of these codons to other amino acids, had been discovered.^[2]^ Today over twenty variations to the standard genetic code^[3]^ have spurred interest in utilizing these variations to encode non-canonical amino acids (ncAAs). Expanding the set of amino acids available for co-translational incorporation by the ribosome opens opportunities to site-specifically introduce new chemistries into proteins and has the potential to transform how we synthesize materials, study protein structure, and understand the translation system.^[4–8]^ For instance, synthesis of high molecular weight, high yielding polypeptides can be achieved inside cells^[9]^ while it can be difficult using other methods.^[10]^

The site-specific incorporation of ncAAs into proteins has been utilized for biophysical studies,^[11,12]^ creating new biocatalysts,^[13]^ synthesizing proteins containing post-translational modifications,^[14–16]^ and understanding translational processes and its evolution over time.^[17]^ Currently over 200 ncAAs have been incorporated into proteins co-translationally.^[18,19]^ These ncAAs include non-canonical α-amino acids (e.g., p-azidophenylalanine, fluorescent amino acids), as well as cyclic and backbone-extended (e.g., β-, γ-, δ-) monomers, among others.^[20–30]^ This is most often achieved in cells by amber suppression whereby the amber stop codon (UAG) is hijacked and used to encode a ncAA through the activity of a ncAA-specific orthogonal translation system (OTS).^[31]^ In general, OTSs are composed of two key components: (i) a suppressor tRNA that has been modified to decode the amber codon (o-tRNA) and (ii) an aminoacyl-tRNA synthetase that has been engineered to specifically recognize an ncAA of interest and aminoacylate it to the o-tRNA (o-aaRS).^[32]^ These components are orthogonal in that they interact principally with an ncAA of interest and have little crosstalk with the native translation components. In practice, these orthogonal components are expressed from a plasmid such as pEVOL,^[33]^ which encodes an o-tRNA as well as two copies of the associated o-aaRS. In the presence of their cognate ncAA, these orthogonal components facilitate the repurposing of amber codons as open channels encoding the ncAA. Historically, a key limitation to OTSs for ncAA incorporation has been premature truncation of the recombinant protein at the UAG codon due to the activity of endogenous release factor-1 (RF-1). During translation, RF-1 recognizes and binds at amber codons, subsequently activating hydrolysis of peptidyl-tRNA to release the peptide chain.^[34]^ Thus, when attempting amber suppression the inherent competition between RF-1 and ncAA-o-tRNAs at amber codons causes inefficient incorporation of ncAAs and premature truncation, reducing the modified protein yield. Within the last decade, this interference was addressed with the completion of an *Escherichia coli* strain lacking all 321 UAG amber stop codons and RF-1, termed *C321.ΔA*. The amber codon is completely orthogonal in this strain and is freed for total dedication to encoding an additional ncAA.^[35]^ Other approaches and strains have also been reported.^[36,37]^ *C321.ΔA* has an increased ability to incorporate multiple ncAAs as compared to other *E. coli* strains, enabling many applications^[15,38–40]^ most notably in biocontainment^[40]^. However, compared to standard commercially available protein production strains, like BL21(DE3), *C321.ΔA* is directly derived from the K-strain *MG1655* (considered wild-type *E. coli*) and has not yet to our knowledge undergone strain development for improved protein production.

In this study, we sought to improve the utility of *C321.ΔA* by introducing genomic mutations and a robust, inducible expression system to enhance the strain’s protein production utility. Given previous successes in improving protein production in *E. coli* systems by removing negative effectors,^[41–43]^ we targeted DNAase *endA*,^[38,44]^ RNAases *rne*^[45]^ and *rnb*,^[46,47]^ and proteases *lon*^[48]^ and *ompT*^[48]^ for functional inactivation in *C321.ΔA*. We show that deletion of nuclease and protease genes in the strain increases production of superfolder green fluorescent protein (sfGFP) containing 2 UAGs by 2.3-and 5.6-fold with p-azidophenylalanine (pAzF) (“click” chemistry; photoreactive crosslinker) and N_6_-(propargyloxycarbonyl)-_L_-Lysine (ProCarb) (pyrrolysine analog), respectively. Next, we introduced the T7 promoter system^[49,50]^ into *C321.ΔA*, given the advantages of high recombinant protein expression using this system. We present the construction of several strains featuring an isopropyl β-_D_-1-thiogalactopyranoside (IPTG) inducible T7RNAP cassette and demonstrate up to a 17-fold improvement in production of sfGFP containing 2 ProCarb residues using these strains. We envision that the library of organisms described here will be an important resource and provide the community with multiple strain options for expression of proteins containing ncAAs with increased protein yield.

## Results and Discussion

### Functional deactivation of proteases and nucleases enhances sfGFP production in C321.ΔA

We hypothesized that reducing the activity of negative effectors of protein synthesis (e.g., proteases and nucleases) would stabilize critical synthetic intermediates and thus increase protein production in *C321.ΔA*. To test this hypothesis, we prepared *C321.ΔA* derivatives featuring combinatorial functional inactivations of two proteases, two RNAses, and one DNAse using multiplex automated genome engineering (MAGE)^[51]^ (**Figure 1A**; **Table 1**). The Lon protease was deactivated using a mutagenic MAGE oligonucleotide to remove its promoter. This mutation is similar to the Lon protease mutation found in *BL21(DE3)* where a transposable element, IS186, inserted directly into the *lon* promoter preventing Lon’s expression.^[52]^ The point mutation D103A was introduced into *ompT* to eliminate proteolytic activity while maintaining its structural motifs in order to preserve OmpT’s possible chaperone function.^[53,54]^ RNAse E, encoded by *rne*, was truncated by inserting a stop codon at nucleotide 131 (*rne131*), a mutation which was previously found to increase mRNA half-life.^[45,55]^ RNAse II (encoded by *rnb* and involved with mRNA degradation^[56]^) and Endonuclease I (encoded by *endA* and capable of generating breaks in double-stranded DNA^[57]^) were similarly truncated by inserting a stop codon followed by a frameshift mutation in the first quarter of their open reading frames.^[51]^ Starting with the parental strain *C321.ΔA*, these mutations were made in single, double, and some triple and quadruple combinations. Mutations were screened by multiplex allele-specific colony (MASC) PCR^[58]^ or colony PCR and confirmed by DNA sequencing. The average doubling times for the MAGE-modified strains were measured in 2xYT media and were determined to be within 12% of the parental strain (**Figure 1B**), suggesting that the gene disruptions did not drastically affect cellular fitness.

**Figure 1.**
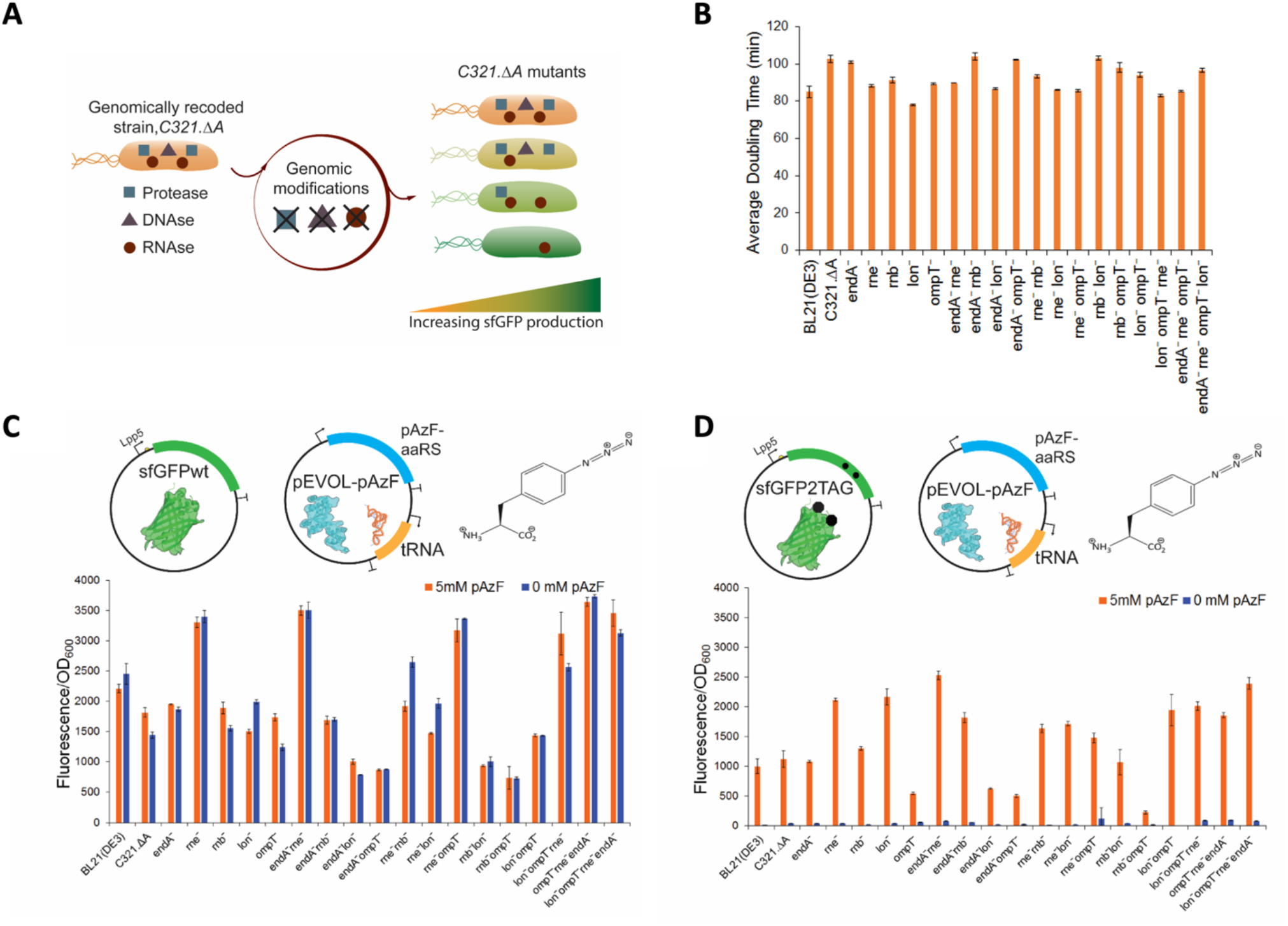
Functional inactivation of nucleases and proteases creates robust, high protein production strains for ncAA incorporation. (A) Mutagenic oligonucleotides were introduced into *C321.ΔA*, targeting two proteases (*lon* and *ompT*), two RNAases (*rne* and *rnb*), and a DNAase (*endA*) for functional inactivation. Through multiple rounds of multiplex automated genome engineering (MAGE) several *C321.ΔA* mutants were generated. (B) *C321.ΔA* mutant strains were grown at 32 °C in 2xYT media in a sterile 96-well polystyrene plates. Optical density at 600 nm was measured every 10 min for 12 h. Error bars represent biological duplicates and technical triplicates. (C) The protein production capability of the modified *C321.ΔA* strains were analyzed by expressing wild-type sfGFP (sfGFP-wt), regulated by a strong endogenous promoter pLpp5, and the pAzF orthogonal translation system expressed on pEVOL-pAzF. For all conditions 1mM IPTG, 0.02% arabinose and 5mM pAzF (orange bars) or 0mM pAzF (blue bars) were added at OD_600_ 0.6-0.8. (D) Modified *C321.ΔA* strains were analyzed for the ability to suppress two amber codons in sfGFP at positions 190 and 212 in the presence (orange) or absence (blue) of 5mM pAzF. For (C) and (D) error bars represent one standard deviation for biological triplicates and technical triplicates.

**Table 1:** Proteases and Nucleases Targeted in this study.

| General Function | Gene | Specific Function | Mutation | Phenotype | Ref. |
| --- | --- | --- | --- | --- | --- |
| DNA Stability | *endA* | Endonuclease I | GGATGT 748 TAACTGA | Truncated at 748 nt | [44] |
| RNA Stability | *rne* | RNase B | GGT 584 TAACTGA | Truncated at 584 nt | [45] |
|  | *rnb* | RNAase E | GACGCC 632 TAACTGA | Truncated at 632 nt | [59] |
| Protein Stability | *lon* | ATP-dependent protease | Removal of promoter | Unknown | [60] |
|  | *ompT* | Outer membrane protease VII | D103A | Elimination of proteolytic activity | [53,54] |

To assess the protein production capacity of *C321.ΔA* and its mutants, we transformed all strains with the pLpp5-sfGFP-wt plasmid, which expresses wild-type sfGFP protein (sfGFP-wt) off a strong IPTG-inducible endogenous promoter, Lpp5^[61]^ along with a pEVOL plasmid encoding the orthogonal translation system (OTS) for *p*-azidophenylalanine (pAzF)^[33]^ (**Figure 1C**). The strains were grown. IPTG was added to induce sfGFP-wt expression, and sfGFP was quantified as a measure of fluorescence/O.D._600_. The *rne*^−^ mutant had the strongest impact on sfGFP-wt expression, implying that mRNA stability may be the largest limitation for expression of sfGFP-wt in *C321.ΔA*. Interestingly, the mutation combination most similar to *BL21(DE3), lon*−*ompT*−, expressed sfGFP-wt at levels 44% less than *BL21(DE3)*. This observation was not explored further, but it most likely stems from the inherent differences between B- and K-strains. Furthermore, we observed that the addition of *endA*− to strains containing *rne*^−^ added a minor boost in sfGFP-wt expression. Ultimately, the top mutants for expression of sfGFP-wt were *rne*^−^,*endA*^−^ *rne*−, and *ompT*^−^ *rne*^−^ *endA*^−^, with the top mutant outproducing both *BL21(DE3)* and the parental strain by 2.6-fold.

We next explored whether the *C321.ΔA* mutant strains could better incorporate ncAAs during protein expression by using an sfGFP-expressing construct containing two amber stop codons (sfGFP-2UAG) (**Figure 1D**). In this case, full length sfGFP expression and fluorescence is dependent on the successful incorporation of pAzF at each amber codon. In the absence of pAzF, any fluorescence measured results from non-specific incorporation at amber codons. Under these conditions, BL21(DE3)’s ability to express sfGFP-2UAG is reduced compared to *C321.ΔA* likely a result of RF-1 being active in *BL21(DE3)*. As was the case with sfGFP-wt, the *rne*^−^ strain and several mutations combined with *rne*^−^ demonstrated increased productivity. The top-performing mutant was *rne*^−^*endA*^−^ with a 2.3-fold improvement as compared to the parental strain. Comparable in performance, *lon*−*ompT*^−^*rne*^−^*endA*^−^ was the next top-performing mutant. Absolute protein expression was quantified by purifying sfGFP using a Strep-tag-Strep-Tactin column after a 20-h expression assay in 1 L 2xYT media (**Table S1**).

In order to test the limits of these systems, top-performing mutants for both sfGFP-wt- and sfGFP-2UAG-expression were tested for the ability to incorporate pAzF at 10 amber codons. We co-transformed top *C321.ΔA* mutants with a plasmid expressing an elastic-like polymer (ELP) containing 10UAGs fused to sfGFP-wt at the C-terminus and the pEVOL-pAzF plasmid, preventing potential loss in fluorescence due to multiple-site ncAA incorporation into sfGFP (**Figure 2A**). In this case, all 10UAGs in the ELP must be suppressed in order for the sfGFP-wt moiety to be expressed and produce fluorescence. The advantage of the genomically recoded strain over *BL21(DE3)*, which contains RF-1, to incorporate multiple ncAAs was especially pronounced here. None of the mutants in this case displayed a significant improvement compared to *C321.ΔA*; however, the system showed a 12-fold improvement over *BL21(DE3)*. This suggests that the ability of the OTS components to incorporate ncAAs, rather than the strain’s protein production capability, is the limiting factor under these conditions.

**Figure 2.**
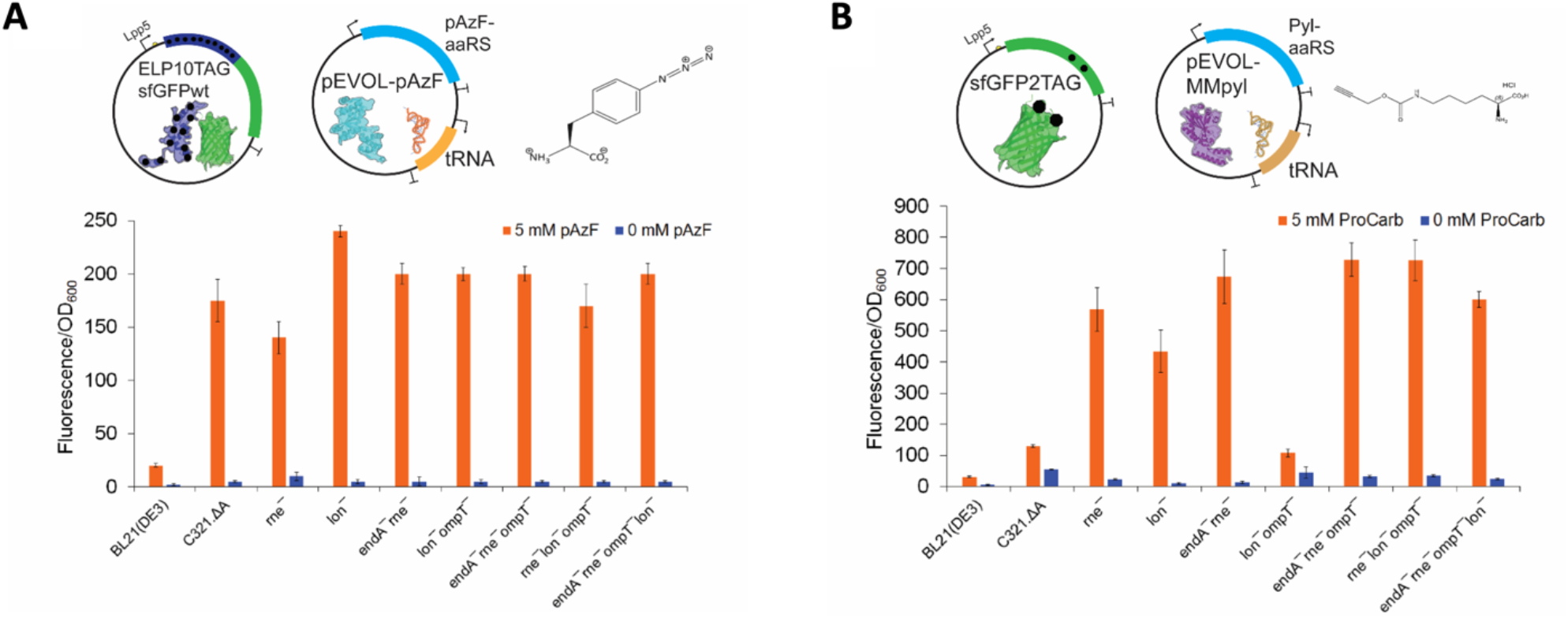
Reducing protease and nuclease activity improves multiple-site ncAA incorporation across orthogonal translation systems. (A) Elastin-like polypeptide (ELP) containing 10 amber codons and fused to sfGFP-wt at its C-terminus was expressed with pEVOL-pAzF in top *C321.ΔA* mutants. For all conditions 1mM IPTG, 0.02% arabinose and 5mM pAzF (orange bars) or 0mM pAzF (blue bars) were added at OD_600_ 0.6-0.8. (B) Expression of sfGFP containing two amber codons for incorporation of the pyrrolysine analog, N_6_- (propargyloxycarbonyl)-_L_-Lysine (ProCarb) using the orthogonal translational system for pyrrolysine was tested in top *C321.ΔA* mutants. For all panels error bars represent one standard deviation for biological triplicates and technical triplicates.

We next tested conditions where mRNA and protein stability may be a limitation by expressing sfGFP2UAG in the presence of a pEVOL plasmid encoding the pyrrolysine (Pyl) OTS system from *Methanosarcina mazei* (pEVOL-MMpyl)^[62]^ (**Figure 2B**). This OTS was chosen because the Pyl synthetase (PylRS) is known to be very difficult to express recombinantly. ^[63,64]^ Because there is no commercial source of Pyl and the ncAA is tedious and expensive to synthesize,^[65]^ we used the pyrrolysine derivative N_6_-(propargyloxycarbonyl)-_L_-Lysine (ProCarb) in these experiments. We observed that the *C321.ΔA* mutants had a drastic improvement over *C321.ΔA* and BL21(DE3) for expression of sfGFP containing two ProCarbs. The top mutant, *endA*−*rne*−*ompT*−, showed a 5.6-fold improvement compared to the parental strain. These results demonstrate the advantage of reducing protease and nuclease activity when using o-aaRSs with known solubility issues such as PylRS. ^[63,64]^

### Genomic introduction of a T7 RNA polymerase cassette increases utility of C321.ΔA

After improving protein production capabilities in *C321.ΔA* we wanted to further enhance its utility by providing transcriptional tuning capabilities. Transcriptional tuning is a powerful tool for efficient recombinant protein production in *E. coli*. Many challenges such as product toxicity, formation of inclusion bodies, and metabolic burden are associated with non-optimal (too high or too low) levels of recombinant protein expression. Tunable expression systems allow for the adjustment of recombinant protein expression using a small molecule inducer to maximally exploit the cell’s metabolic capability. Thus, the ability to tune recombinant protein expression is a staple for many protein expression projects. Within this realm, use of the T7 RNA polymerase (T7RNAP) within *BL21(DE3)* is the most popular approach for producing proteins due to the enzyme’s high activity, tunability, and orthogonality. To use this system, a gene of interest is cloned behind a T7 promoter and recognized exclusively by the phage T7RNAP encoded on the genomic DE3 cassette and induced by the addition of IPTG.^[48]^ This allows for highly productive and orthogonal recombinant protein expression that is tunable by controlled addition of IPTG. To leverage the power of the T7RNAP in *C321.ΔA*, a synthetic T7RNAP cassette was synthesized by amplifying T7RNAP from the BL21(DE3) genome and adding an upstream terminator (to transcriptionally isolate the cassette), a CmR gene as a selectable marker, and 45 bp of genomic homology to the genomic insertion site on the 5’ and 3’ end to facilitate incorporation into the genome (**Figure S1**). Using λ-red mediated homologous recombination,^[58,66]^ the cassette was inserted into the top *C321.ΔA* mutant strains. Using MAGE, the CmR marker was removed, and finally the full cassette and insertion site were verified by sequencing to yield a series of *C321.ΔA-T7 strains* (**Figure 3A**).

**Figure 3:**
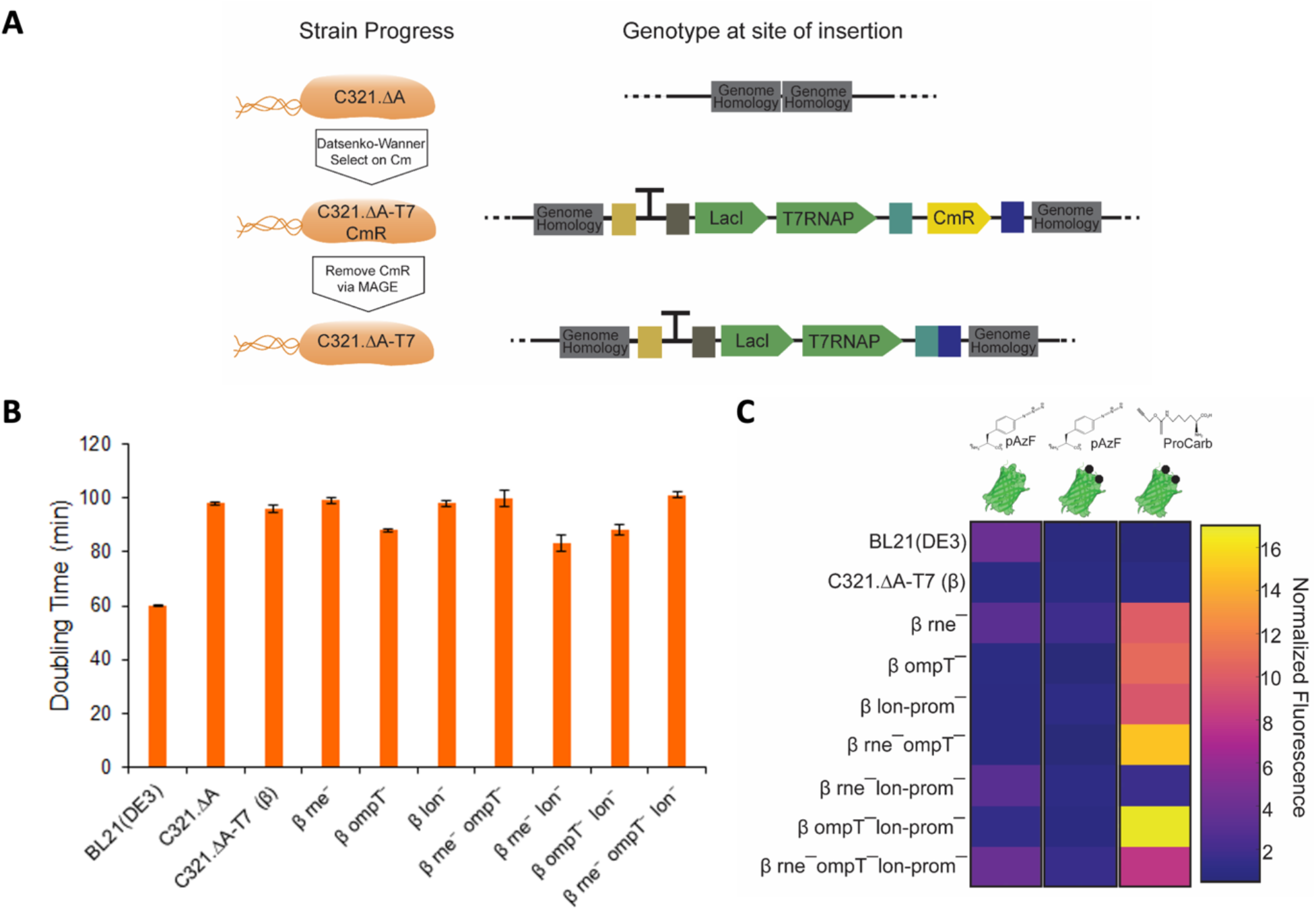
Genomic insertion of the T7RNAP cassette improves utility of *C321.ΔA* mutant strains. (A) The T7RNAP cassette was inserted into top *C321.ΔA* mutant strains using Datsenko- Wanner. After recovery, transformed cells were plated on LB plates containing 34 μg/mL chloramphenicol, selecting for strains that incorporated the T7RNAP cassette. The selectable marker was then removed through several cycles of multiplex automated genome engineering (MAGE). The resultant strains were termed *C321.ΔA-T7*. (B) Strains were grown at 32 °C in 2xYT media in a sterile 96-well polystyrene plates. Optical density at 600 nm was measured every 10 min for 12 h. Error bars represent biological duplicates and technical triplicates. (C) Heat map depicts the normalized fluorescence (Fluorescence/OD_600_) for various reporter proteins relative to *C321.ΔA-T7(β)* within each vertical condition. Column 1: pET28a-sfGFP-wt + pEVOLpAzF + 5 mM pAzF + 1 mM IPTG + 0.02% Arabinose; Column 2: pET28a-sfGFP2UAG + pEVOLpAzF + 5 mM pAzF + 1 mM IPTG + 0.02% Arabinose; Column 3: pET28a-sfGFP2UAG + pEVOL-MMpyl + 5 mM ProCarb + 1 mM IPTG + 0.02% Arabinose. Normalized Fluorescence data is shown in **Figure S4** and **S5**.

To test functionality of the T7RNAP cassette, a reporter plasmid expressing sfGFP-wt or sfGFP-2UAG regulated by a T7 promoter was transformed into the *C321.ΔA-T7* strains along with pEVOL-pAzF and the ability of the strains to express sfGFP using the T7RNAP was assessed. Here, all strains including BL21(DE3) expressed the reporter proteins at levels much lower than sfGFP-wt/2UAG expression regulated by Lpp5 (**Figure S2**). Because T7 systems often utilize pET plasmids, the reporter plasmids were switched to a pET28a backbone.^[67]^ With the new plasmids BL21(DE3)’s expression of sfGFP-wt (**Figure S3A**) increased to near the same levels as expression of pLpp5-sfGFPwt. However, expression of sfGFP-wt, directed by T7RNAP, remained low in the *C321.ΔA* mutants.

We hypothesized that T7RNAP-based expression in *C321.ΔA* could be improved by introducing genomic mutations similar to those present in BL21 (DE3) and its derivatives. To test this, we reconstructed the *C321.ΔA* strains containing T7RNAP by addition of the T7RNAP cassette into *C321.ΔA* (to yield strain *β*), followed by the combinatorial introduction of inactivating mutations to *rne, ompT*, and *lon*. The doubling times of the resulting strains in 2xYT media at 32 °C were within 18% of the parental strain (**Figure 3B**). When assessing T7RNAP-driven expression using these strains, we observed an increase in productivity of 2.2-fold and 3.1-fold for sfGFP-wt/pAzF and sfGFP-2UAG/pAzF respectively (**Figure S4**).

In particular, *β rne*−*ompT*−*lon*−, featuring the same combination of mutations as BL21 (DE3), showed an improvement in sfGFP-wt expression of 3.7-fold over the parental strain (**Figure 3C**). When expressing pET28a-sfGFP2UAG with pEVOL-pAzF, *β rne*^−^ showed a 1.9-fold improvement over *β* and 2.3-fold improvement over BL21(DE3). It appears that no matter the RNA polymerase used for expression, *rne*^−^ is the most beneficial mutation for expression of sfGFP in *C321.ΔA* strains. Lastly, when expressing sfGFP-2UAG with pEVOL-MMpyl, we observed a 17-fold improvement compared to *β*. In this case, we suspect the highest fold improvement was observed due to the poor solubility of PylRS.

### Conclusions

The recoded strain *C321.ΔA* has a fully orthogonal amber codon for site-specific ncAA incorporation. However, it has not previously been optimized for protein production *in vivo*. To address this gap, we applied MAGE^[51]^ to generate a series of *C321.ΔA* derivatives with combinatorial knockouts of several nucleases and proteases. Protein yield from each strain was quantified as a function of sfGFP fluorescence/O.D._600_ in order to assess the impacts of these knockouts on protein production. This analysis revealed that the functional inactivation of three targets (*ompT*−*rne*−*endA*−) improved sfGFPwt production by ∼2.6-fold, while inactivation of *rne* and *endA* improved the production of sfGFP featuring two pAzF residues (sfGFP2pAzF) by ∼2.3- fold. Pushing the limits of the system, inactivation of the *Lon* protease dramatically improved expression of elastin-like polymer featuring 10 pAzF residues. Finally, use of the pyrrolysyl-tRNA synthetase improved ∼5.6-fold in a top mutant as compared to the parental strain, suggesting applicability of our approach to cases in which mRNA and/or protein stability are known issues.^[63,64]^ Notably, across all trials and experiments the functional inactivation of the RNAse *rne* was the most impactful.

To introduce precise control of target protein expression via transcriptional tuning, we applied λ-Red-mediated homologous incorporation to introduce a synthetic cassette encoding an IPTG-inducible T7 RNA polymerase into our strains. When this cassette was inserted into *C321.ΔA* prior to the combinatorial inactivation of *ompT, rne*, and *lon*, improvements of up to 3.7- and 1.9-fold were observed for sfGFPwt and sfGFP2pAzF, respectively, as compared to the parental strain. Incorporation of pyrrolysine also improved significantly in these strains, again likely as a result of improved PylRS solubility. Importantly, the top-performing T7 strains developed in this study all significantly outperformed BL21 (DE3) in terms of both yield and ability to incorporate ncAAs at amber codons.

Looking forward, as our ability to increase efficiencies of OTSs improve so will the need for optimized strains for the production of proteins containing ncAAs. For example, new efforts to engineer tethered ribosomes in cells offer exciting new dimensions for expanding the chemistry of life.^[68–72]^ For example, Orelle et al. first demonstrated the evolvability of tethered ribosomes by selecting otherwise dominantly lethal rRNA mutations in the peptidyl transferase center that facilitate the translation of problematic protein sequences.^[71]^ This, supported by increases in biomanufacturing should make possible new avenues in engineering molecular translation systems.

## Materials and Methods

### Reagents, Buffers and plasmids

Chemicals and media were purchased from Sigma Aldrich (St. Louise, MO, USA) unless otherwise designated. Phusion High-Fidelity DNA Polymerase, Taq DNA polymerase with Standard Taq Buffer, T4 DNA ligase, dNTP, Quick-load DNA Ladders, BL21(DE3) and restriction endonuclease were purchased from New England Biolabs (NEB, Ipswich, MA, USA). Multipex PCR Kits used for MASC PCR were purchased from QIAGEN (Hilden, NRW, DE). Plasmids were extracted using Omega E.Z.N.A DNA Isolation Kit (Omega Bio-Tek, Norcross, GA, USA). DNA was column purified or gel extracted using OMEGA HiBind DNA Mini Columns and OMEGA E.Z.N.A Gel Extraction Kit, respectively. Genomic DNA was isolated with Omega E.Z.N.A. Bacterial DNA Kit. All DNA oligonucleotides were purchased from Integrated DNA Technologies (IDT, Coralville, IA, USA). The ncAA pAzF was purchased from P212121, LLC (Ann Arbor, MI, USA) and ProCarb was purchased from BioFine, Inc (Vancouver, BC, CA). SYBR Safe, used in all agarose gels, and DH5α were purchased from Thermo Fisher Scientific (Waltham, MA, USA). Synthetic *E. coli* C321.ΔA (GenBank: CP06698.1) was received as a gift from Farren Isaacs. All oligonucleotides used for cloning are shown in **Table S2**. All vectors were cloned using Gibson Assembly.^[73]^ pLpp5 plasmids were derived from pDTT1 vector.^[74]^ pET vectors were derived from pET28a vectors.

### Construction of C321.ΔA mutants

The strains in this study were generated from *C321.ΔA*^[75]^ by disrupting genes of interest using mutagenic oligonucleotides via MAGE^[58]^ (**Table S2**). Cultures were grown in LB-Lennox media (10 g/L Trypton, 5 g/L Yeast Extract, and 5 g/L NaCl) at 32°C and 250 rpm throughout the MAGE cycle steps.^[58]^ Single, double, several triple and quadruple mutations were made to endA, rne, rnb, lon, and ompT, to investigate the effect of reduced nuclease and protease activity on expression of protein containing multiple ncAAs. Multiplex allele-specific colony (MASC) PCR was performed to screen for gene mutations by using wild-type forward (-wt-f) or mutant forward (- mut-f) primers and reverse primers (-r; **Table S2**). Wild-type and mutant forward primers were identical except at the 3’-ends of the oligonucleotide, and the reverse primers were used for detection of both wild-type and mutant alleles. The mutant allele was amplified using the mutant forward and reverse promoter set (-mut-f and –r) which resulted in a band on an electrophoresis gel but not with the wild-type forward and reverse primer set (-wt-f and –r). MASC PCR was performed in 10 μL reactions by using a Multiplex Master Mix at 95°C for 15 min, with 30 cycles of 95°C for 30 s, 65°C for 30 s, and 72°C for 1 min, and a final extension of 72°C for 5 min. Selection for *lon* mutants were performed separately in 10 μL reactions using Taq DNA polymerase with Standard Taq Buffer at 95°C for 15 min, with 20 cycles of 95°C for 30 s, 55°C for 30 s, and 68°C for 2min, and a final extension of 68°C for 5 min. Mutant alleles were screened by running PCR products on a 2% agarose gel and confirmed by DNA sequencing by using sequencing primers (**Table S2**).

### Growth Curves

Overnight cultures of strains were grown in 2X YT (16 g/L Trypton, 10 g/L Yeast Extract, and 5 g/L NaCl) media at 32°C at 250rpm and were diluted 1:50 in 100 μL of 2X YT media. Diluted cultures (100 μL) were added to 96-well polystyrene plates (Costar 3370; Corning Incorporated, Corning, NY, USA). The OD_600_ was measured at 10 min intervals for 20 hr at 32°C in orbital shaking mode on a SynergyH1 plate reader (Biotek, Winooski, VT, USA). Growth data for each strain was obtained from three replicate wells and three independent cultures. Doubling time was calculated during exponential growth phase.

### Assaying expression of GFP

Strains were freshly transformed with the plasmids of interest. A single colony was inoculated into 5 mL of 2X YT media with 35 μg/mL Kanamycin and 25 μg/mL Chloramphenicol (Kan_35_Cm_25_) grown overnight at 32°C, 250 rpm. Overnight cultures were diluted 1:50 into 5 mL of fresh 2X YT media Kan_35_Cm_25_ in triplicate and grown at 32°C at 250 rpm. OD_600_ was monitored on a Libra S4 spectrophotometer (Biochrom, Cambridge, UK) until OD_600_ 0.6-0.8 at which point cultures were induced. Inducers consisted of either 5 mM ncAA, 1 mM IPTG, and 0.02% arabinose or 0 mM ncAA, 1 mM IPTG, and 0.02% arabinose. Cultures were allowed to express for 20-24 hrs after induction prior to harvest. To assay fluorescence, overnight cultures were diluted 10-fold in 2X YT media Kan_35_Cm_25_. The OD_600_ of the 10-fold dilution was measured on a NanoDrop 2000c (Thermo Scientific, Waltham, USA) and multiplied by ten. 100 μL of the 10-fold dilution was added to 96-well polystyrene plates (Costar 3603) in triplicate. Fluorescence of the plates were measured on a Synergy H1 plate reader with a gain of 60. Normalized fluorescence was obtained by dividing fluorescence reading (normalized to 2X YT media Kan_35_Cm_25_ wells) by OD_600_ read on the NanoDrop 2000c.

### Construction of T7RNAP cassette

The T7RNAP cassette was assembled from three pieces: a terminator (TM) piece, a T7RNAP piece, a CmR piece (**Table S3**). To transcriptionally isolate from the cassette a 5’ terminator was designed upstream the T7RNAP piece. The strong synthetic terminator (L3S2P21)^[76]^ was selected to avoid potential homology with native terminators during genomic insertion. The terminator was order was from IDT as a sense and antisense oligonucleotide (**Table S2**). The T7RNAP part was amplified from *BL21(DE3)* genomic DNA. The T7RNAP PCR was performed using Phusion with EDC408 and EDC323 primers, 5 ng genomic DNA per μL of PCR reaction, 3% DMSO at 98°C for 15 min, with 30 cycles of 98°C for 30 s, 55°C for 30 s, and 72°C for 3 min, and a final extension of 72°C for 25 min. The CmR piece PCR was performed using Phusion with EDC413 and EDC414 primers and the pAM552C plasmid^[71]^ at 98°C for 15 min, with 30 cycles of 98°C for 60 s, 55°C for 30 s, and 72°C for 45 s, and a final extension of 72°C for 25 min. The T7RNAP and CmR PCR reactions each received 1 μL of DpnI per 20 μL of PCR reaction and were incubated at 37°C for 2 hr. The PCR reactions were column purified and run on a 0.7% agarose gel at 90 V for 45 min. The correct sized band was cut out of the gel and column purified. All three parts were then pool together at equal molar concentrations (75 ng of DNA total) in an overlap PCR reaction using Phusion, 3% DMSO at 98°C for 10 min, with 15 cycles of 98°C for 30 s, 55°C for 30 s, and 72°C for 4 min, and a final extension of 72°C for 10 min. The overlap PCR was then diluted 20-fold into a second PCR reaction with EDC410 and EDC414 primers at 98°C for 3 min, with 24 cycles of 98°C for 30 s, 55°C for 30 s, and 72°C for 4 min, and a final extension of 72°C for 10 min. PCR reactions were then column purified and run on a 0.7% agarose gel at 90 V for 45 min. The correct sized bands were cut out and column purified. Next, 45 bp of genomic insertion site homology was added to the 5’ and 3’ end of the assembled T7RNAP cassette using Phusion, 3% DMSO with JGP139 and JGP140 primers at 98°C for 3 min, with 25 cycles of 98°C for 60 s, 65°C for 30 s, and 72°C for 7 min, and a final extension of 72°C for 10 min. PCR reactions were column purified, run on a 0.7% agarose gel at 90 V for 45 min. The correct sized bands were cut out and column purified. The sequence of the fully assembled cassette was confirmed via sequencing.

### Datsenko-Wanner of T7RNAP cassette

The T7RNAP cassette was inserted using the λ-red homologous recombination method for PCR products.^[58,66]^ The *C321.ΔA* strain contains the λ-red recombinase machinery on its genome which enables quick modification of the genome without scars. Briefly, 3 mL of LB-L media was inoculated with overnight culture of the strain of interest at a 1:50 dilution. Cultures were grown at 32°C, 250 rpm until OD_600_ reached 0.7 as read on a Libra S4. The culture was heat shocked at 42°C for 15 min at 100 rpm to activate expression of the λ-red recombinase machinery. Cultures were place on ice for at least 15 min to cool cells down, spinning the culture tube in ice every 3 min. Next, 1 mL of culture was harvested and washed twice with ice-cold sterile deionized water, pelleting cells at 13,000 g at 4°C. The cell pellets were resuspended with 10 ng of the T7RNAP cassette in 100 μL of ice-cold sterile deionized water and electroporated. Cells were then recovered in 1 mL LB-L for at least 3 hr at 32°C, 250 rpm and plated on Cm_34_ plates for 1-3 days at 30°C.

### Screening for full T7RNAP cassette insertion

Cells that genomically inserted the CmR portion of the cassette grew on the Cm_34_ plates. To screen for full insertion of the cassette colony PCR was performed. Colonies on the Cm_34_ plate were picked and inoculated into 100 μL LB-L Cm_25_ media in 96-well polystyrene plates (Costar 3370) incubated at 32°C, 250 rpm for at least 3 hr. The cultures were used as the template in colony PCR reactions. To screen for 5’ portion of T7RNAP a PCR reaction was performed with MASC PCR reactions using JGP173 and JGP292 primers at 95°C for 15 min, with 30 cycles of 95°C for 30s, 52°C for 30 s, and 72°C for 1 min, and a final extension of 72°C for 10 min. PCR reactions were run on a 2% gel, 110V 45 min. Colony PCR was repeated at a larger scale for the colonies that resulted in a band, reactions were column purified and submitted for sequencing using JGP173, EDC280 and JGP292 primers. Positive sequence hits were screened for the full T7RNAP region being inserted using Multiplex Master Mix with EDC282 and JGP153 primers at 95°C for 15 min, with 30 cycles of 95°C for 30 s, 53°C for 30 s, and 72°C for 1.5 min, and a final extension of 72°C for 10 min. The PCR reaction were run on a 2% agarose gel. Colony PCR was repeated at a larger scale for the colonies that resulted in a band. The reactions were column purified and submitted for sequencing using EDC282, EDC283, EDC284, EDC285, and JGP153 primers.

### Removing antibiotic resistance marker

Clones with full T7RNAP cassette present then underwent MAGE to remove the CmR gene using a mutagenic oligonucleotide, JGP389, with homology on the 5’ and 3’ end of the CmR gene. After 8 cycles of MAGE, overnight cultures were plated on LB plates at 10^−6^ dilutions in LB-L Cb_50_ media. Colonies were replica-plated onto LB-Cb_100_ and LB Cb_100_Cm_34_ plates and incubated at 32°C overnight. Colonies that grew on LB-Cb_100_ plates and not LB-Cb_100_Cm_34_ plates underwent PCR using Multiplex Master Mix with EDC413 and JGP211 primers at 95°C for 15 min, with 30 cycles of 95°C for 30 s, 54°C for 30 s, and 72°C for 1.5 min, and a final extension of 72°C for 10 min. For positive hits colonies the PCR reactions were repeated at a larger scale, column purified and submitted to sequencing with EDC413 and JGP211 primers to confirm the CmR gene was completely removed.

### Full-length sfGFP purification and quantification

Strains were freshly transformed with the plasmids of interest. A single colony was inoculated into 5 mL of 2X YT media with Kan_35_Cm_25_ and grown overnight at 32°C at 250 rpm. Overnight cultures were diluted 1:50 into 40 mL of fresh 2X YT media Kan_35_Cm_25_ and grown at 32°C, 250 rpm. OD_600_ was monitored on a NanoDrop 2000c until OD_600_ 0.6-0.8 at which point cultures were induced with 5 mM ncAA, 1 mM IPTG, and 0.02% arabinose. Cultures were harvested after 20 hr after induction by pelleting 30 mL of culture at 5,000 g for 10 min at 4°C. The pellet was resuspended in 0.8 mL of 1X phosphate-buffered saline (PBS) buffer for every 1 g of wet cell pellet. Cells were lysed at a frequency of 20 kHz and an amplitude of 50% using a Q125 Sonicator (Qsonica, Newton, CT, USA) with a 3.75 mm diameter probe^[77]^ for 5 cycles of 45 s sonication and 59 s sitting on ice. The input energy (Joules) per cycle averaged to 274. Lysed samples were then centrifuged at 21,000 rpm for 10 min at 4°C. Supernatant was collected as the soluble fraction. Full-length sfGFP was purified from the soluble fraction by using a C-terminal strep-tag and 0.2 mL gratify-flow Strep-Tactin Sepharose mini-columns (IBA GmbH, Gottingen, DEU). Purified sfGFP was measured using a Quick Start Bradford Kit (BioRad, Hercules, CA, USA) in biological triplicate and technical triplicate.

## Associated Content

The Supplemental Information:

Supplemental Table 1. Quantification of *in vivo* protein concentrations of top mutant strains; Supplemental Figure 1. Construction of the T7RNAP cassette; Supplemental Figure 2. Expression of sfGFP in top *C321.ΔA* mutants containing genomically expressed T7RNAP cassette; Supplemental Figure 3. Expression of sfGFP in *C321.ΔA-T7* strains using a pET reporter plasmid; Supplemental Figure 4. Expression of sfGFP in *C321.ΔA-T7* strains produced by first inserting T7RNAP, followed by the introduction of protein and nuclease mutations; Supplemental Figure 5. Expression of sfGFP-2UAG utilizing the pyrrolysine orthogonal translation system in β strains; Supplemental Table 2. Oligonucleotides used in this study; Supplemental Table 3. T7RNAP cassette parts.

## Abbreviations

(T7RNAP): T7 RNA polymerase
(ProCarb): N_6_-(propargyloxycarbonyl)-_L_-Lysine
(OTS): orthogonal translation system
(o-aaRS): orthogonal aminoacyl-tRNA synthetase
(o-tRNA): orthogonal tRNA
(RF-1): release factor-1
(IPTG): isopropyl β-_D_-1-thiogalactopyranoside
(pAzF): p-azidophenylalanine
(MAGE): multiplex automated genome engineering
(ELP): elastic-like polymer
(Pyl): pyrrolysine
(*M. mazei*): *Methanosarcina mazei*
(PylRS): pyrrolysine synthetase.

## Author Information

J.G.P and E.D.C constructed the strains. E.D.C designed the T7RNAP cassette. J.G.P, O.W. and E.D.C. constructed, introduced and verified T7RNAP genomic insertion. J.G.P, E.D.C and M.C.J. designed experiments. J.G.P. and E.D.C. performed the experiments. J.G.P. and M.C.J. wrote the manuscript. All authors discussed the results and commented on the manuscript. M.C.J. has a financial interest in Pearl Bio. M.C.J.’s interests are reviewed and managed by Northwestern University in accordance with their conflict of interest policies. All other authors declare no conflicts of interest.

## Acknowledgements

This work was supported by the National Science Foundation (NSF) (MCB-1716766), the Office of Naval Research (N00014-11-1-0363), the Army Research Office (W911NF-20-1-0195, W911NF-18-1-0200), the Human Frontiers Science Program. RGP0015/2017, and the David and Lucile Packard Foundation (2011-37152), and the NIH Initiative for Maximizing Student Development (IMSD) (PAR-17-053). The authors would like to thank Professor Joshua Leonard for his critical review of the manuscript and Professor Peter G. Schultz for providing the pEVOL plasmid.

## Supplementary Information for

**Supplemental Table 1.**
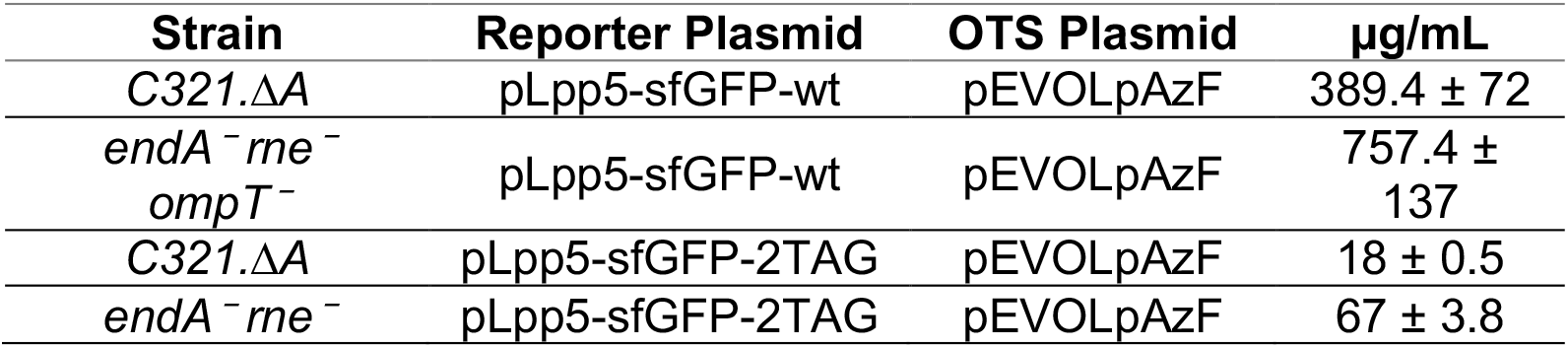
Quantification of *in vivo* protein concentrations of top mutant strains.

| Strain | Reporter Plasmid | OTS Plasmid | µg/mL |
| --- | --- | --- | --- |
| C321.ΔA | pLpp5-sfGFP-wt | pEVOLpAzF | 389.4 ± 72 |
| *endA*<sup>-</sup>*rne*<sup>-</sup><br>*ompT*<sup>-</sup> | pLpp5-sfGFP-wt | pEVOLpAzF | 757.4 ±<br>137 |
| C321.ΔA | pLpp5-sfGFP-2TAG | pEVOLpAzF | 18 ± 0.5 |
| *endA*<sup>-</sup>*rne*<sup>-</sup> | pLpp5-sfGFP-2TAG | pEVOLpAzF | 67 ± 3.8 |

**Supplemental Table 2.**
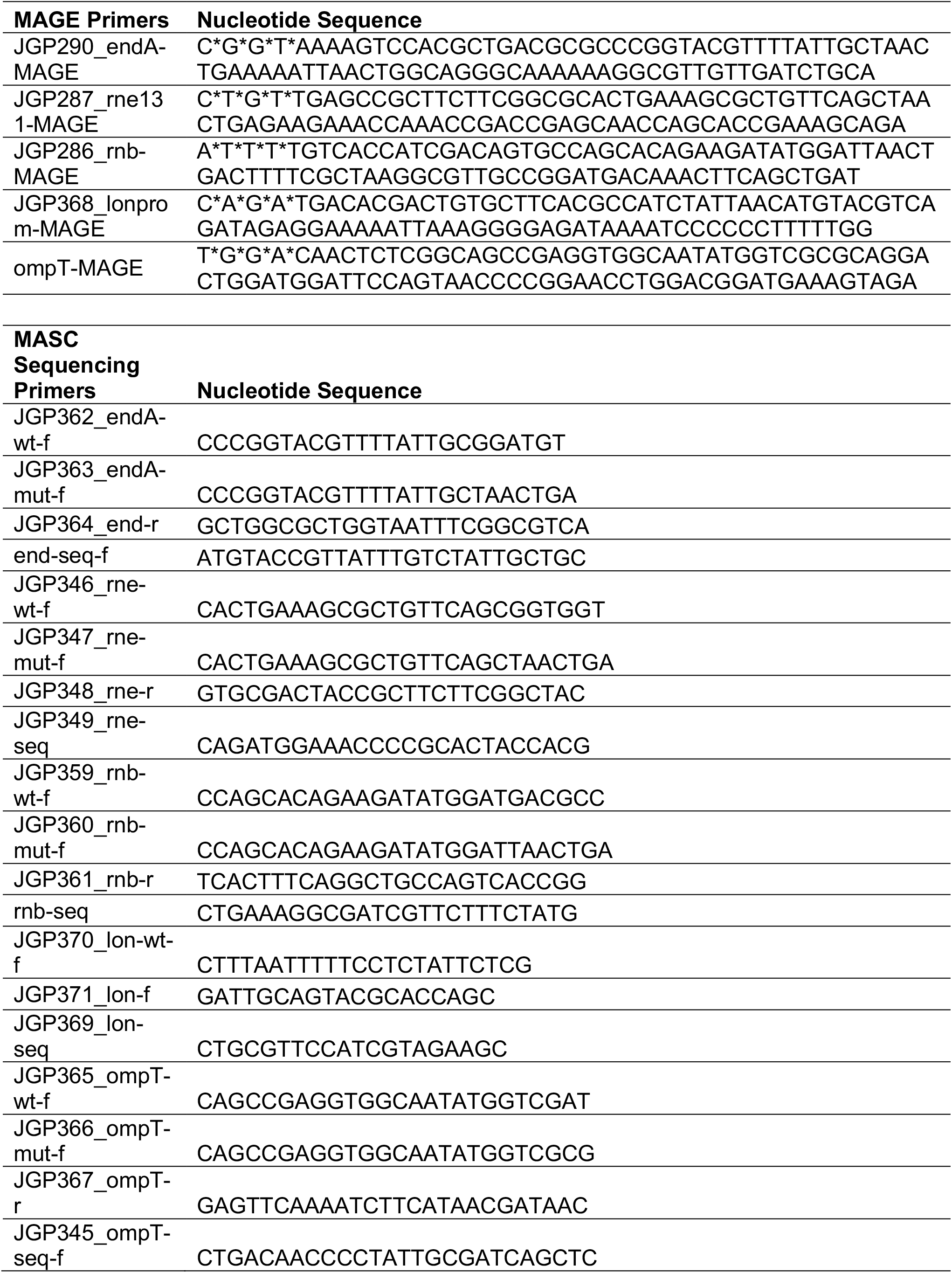
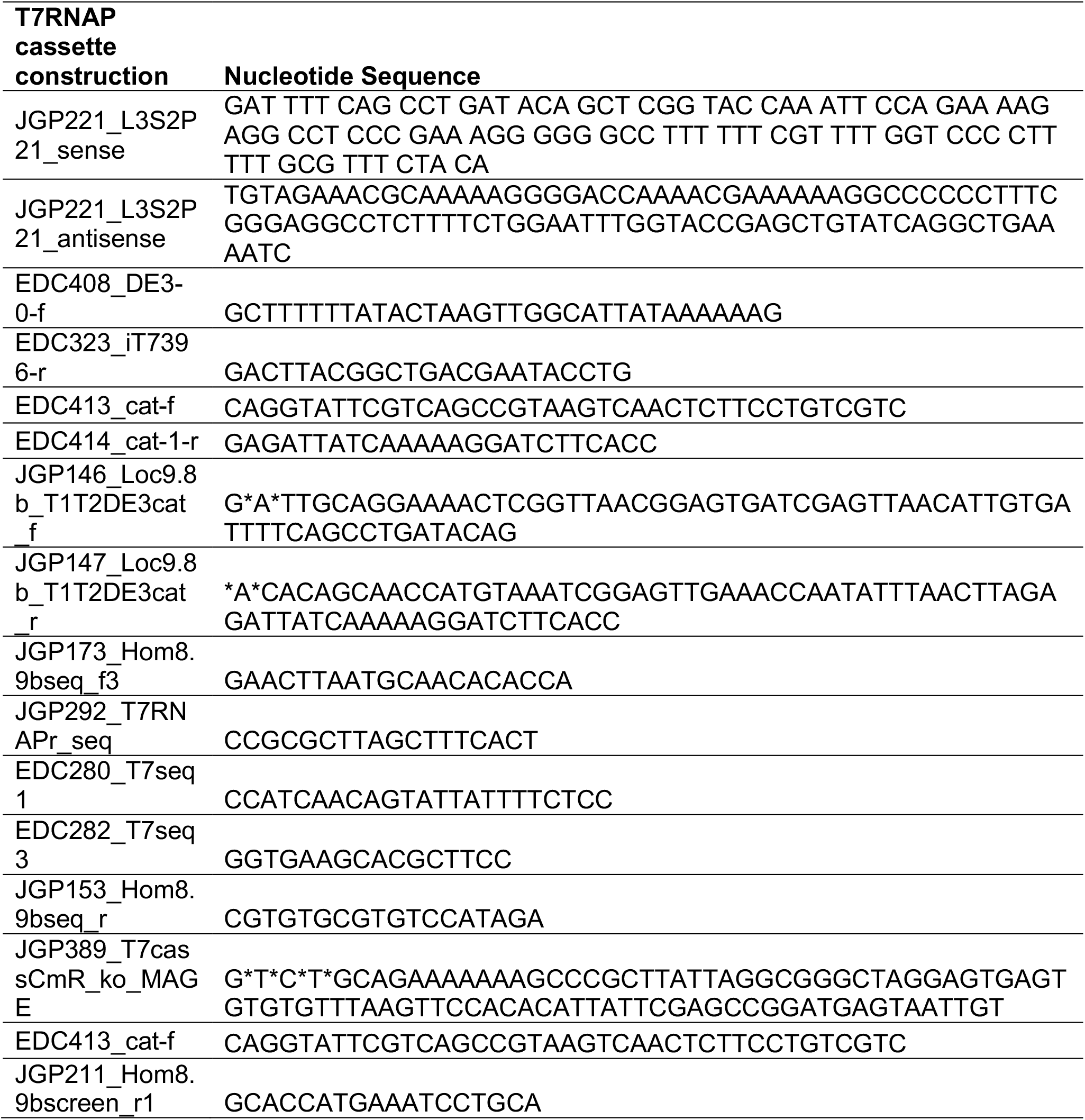
Oligonucleotides used in this study.

**Supplemental Table 3.**
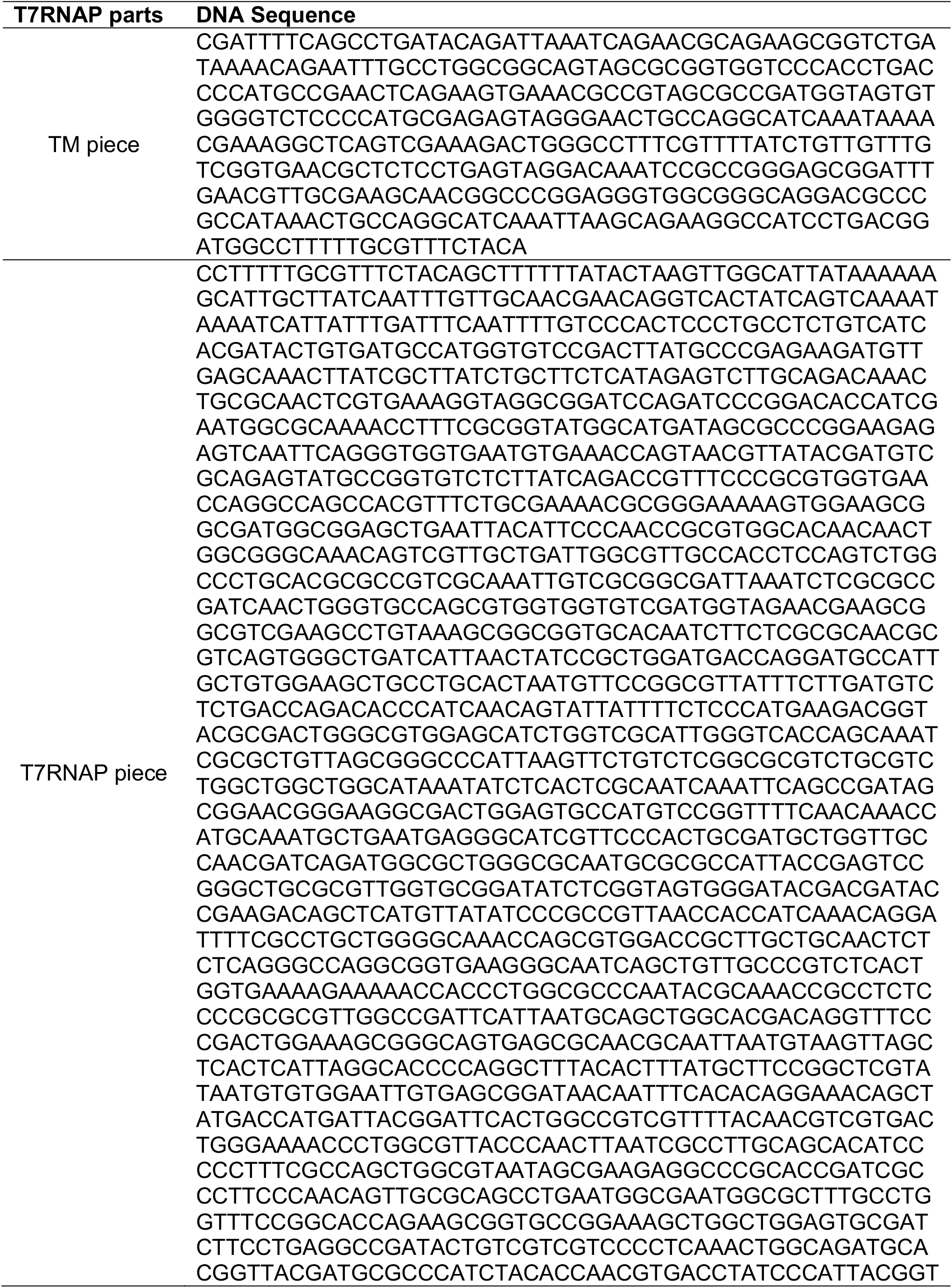
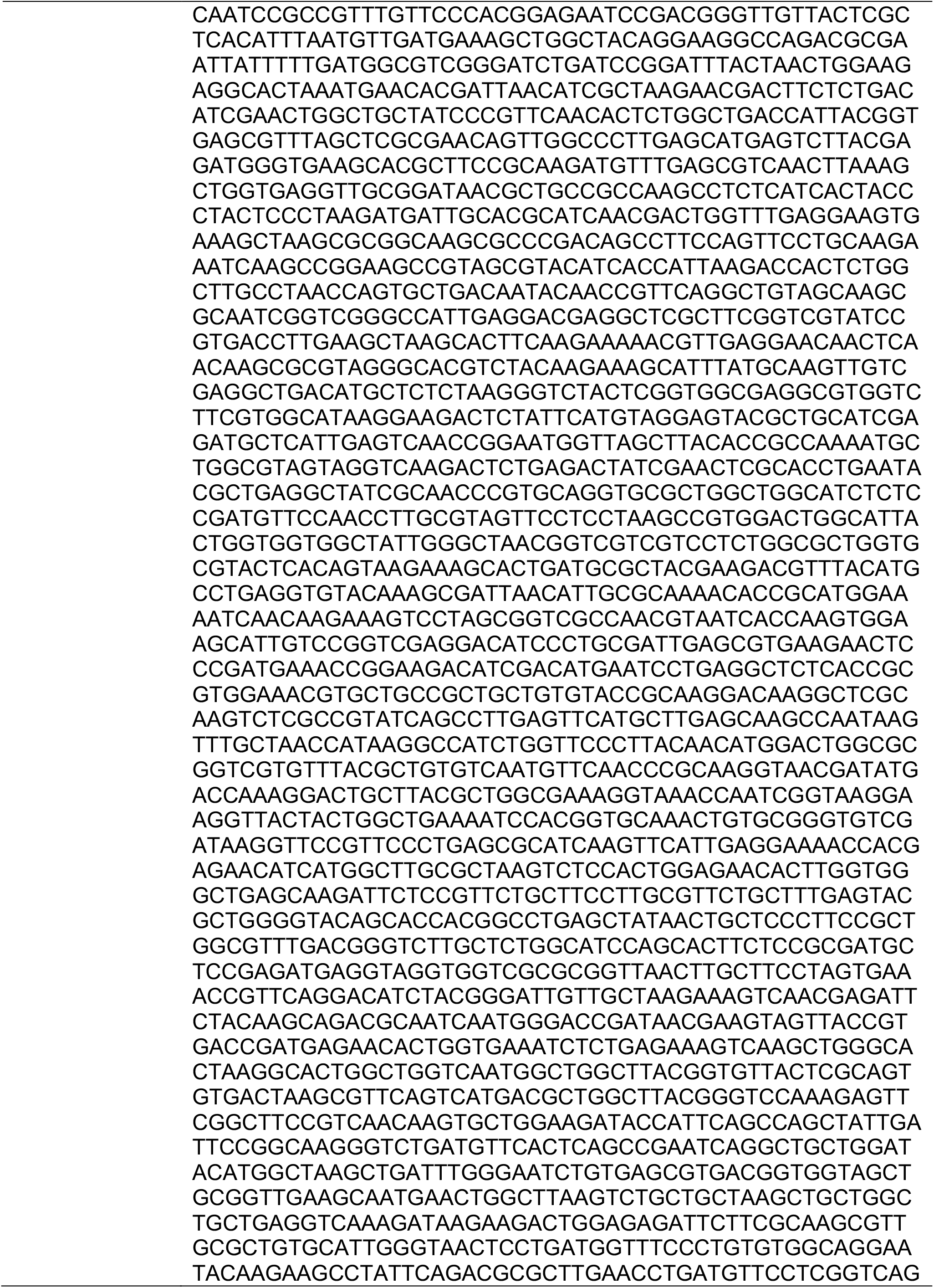
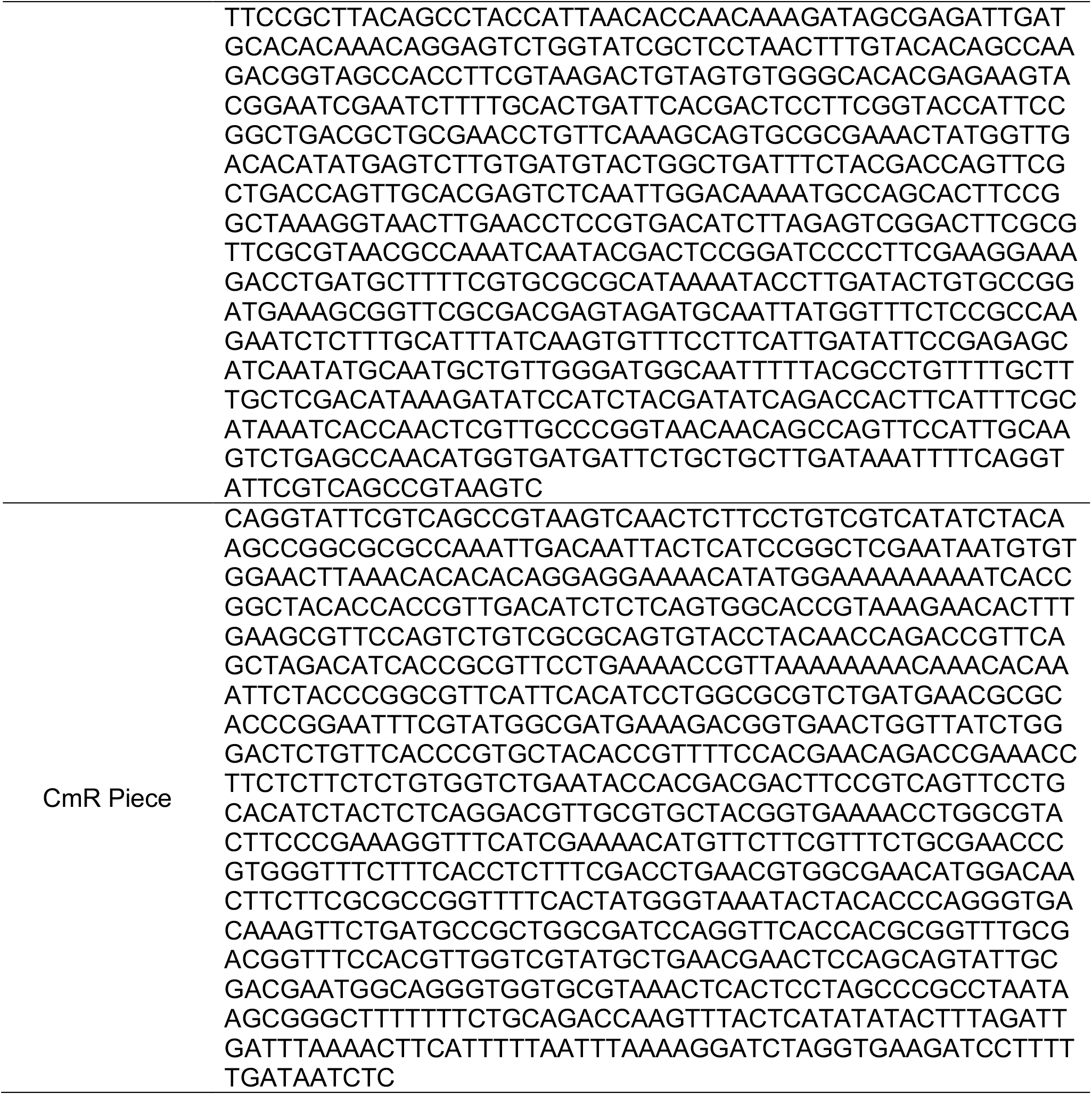
T7RNAP cassette parts.

**Supplemental Figure 1.**
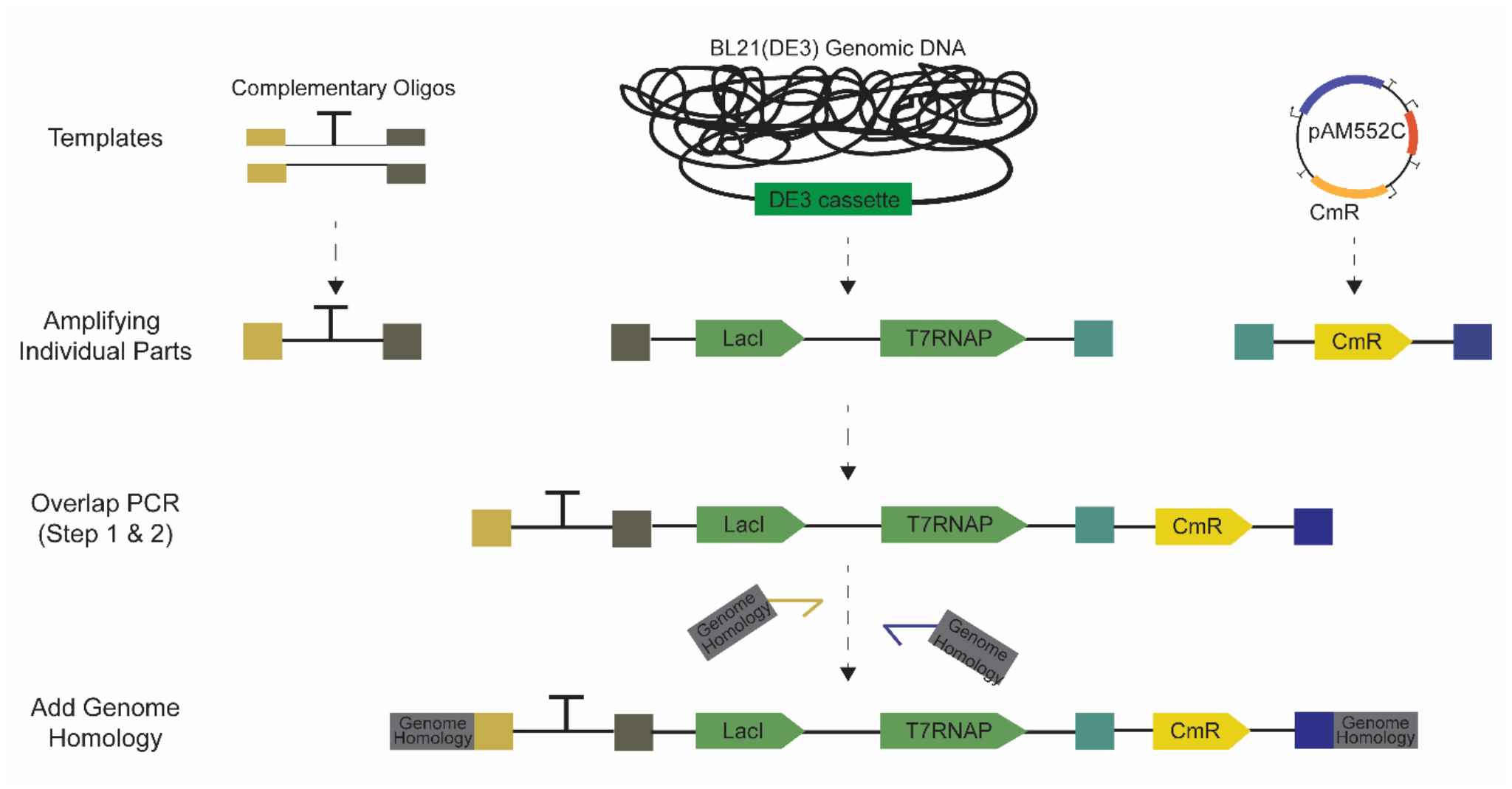
Construction of the T7RNAP cassette. The T7RNAP cassette consists of three parts: 5’ Terminator (TM), T7 RNA Polymerase (T7RNAP) and antibiotic resistance marker CmR. These parts were amplified from various sources and were stitched together via overlap PCR using a two-step amplification process. During the stitching process homology to the genomic insertion site was added to the 5’ and 3’ end of the cassette.

**Supplemental Figure 2.**
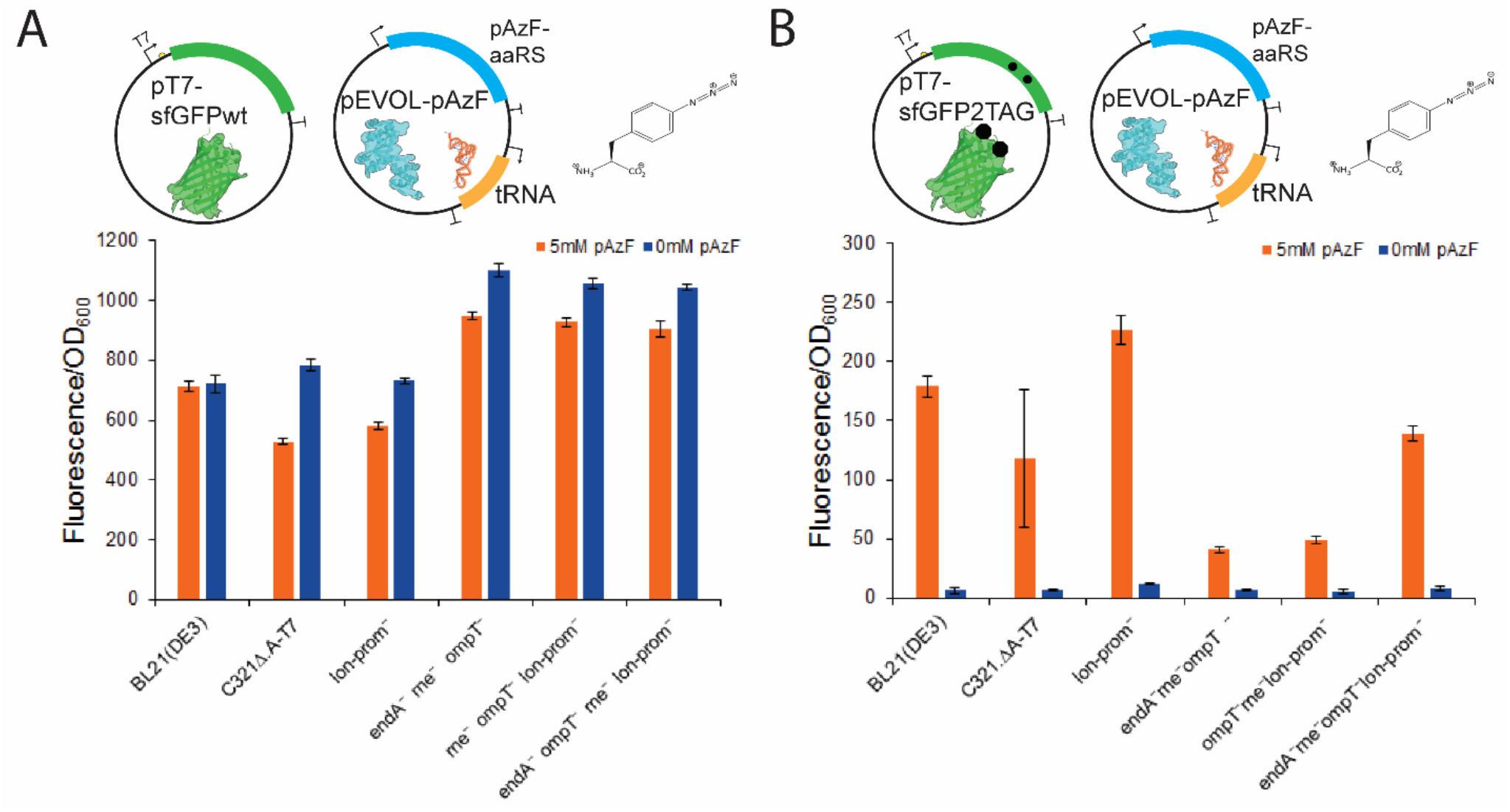
Expression of sfGFP in top *C321.ΔA* mutants containing genomically expressed T7RNAP cassette. A) T7RNAP function was tested in the top *C321.ΔA* mutants by expressing wild-type sfGFP (sfGFP-wt) with the pAzF orthogonal translation system expressed on pEVOL-pAzF. For all conditions 1 mM IPTG, 0.02% arabinose and 5 mM pAzF (orange bars) or 0mM pAzF (blue bars) was added at OD_600_ 0.6-0.8. B) Modified *C321.ΔA-T7* strains were analyzed for the ability to suppress two amber codons at positions 190 and 212 in the presence (orange) or absence (blue) of 5 mM pAzF. For all panels error bars represent one standard deviation for biological triplicates and technical triplicates.

**Supplemental Figure 3.**
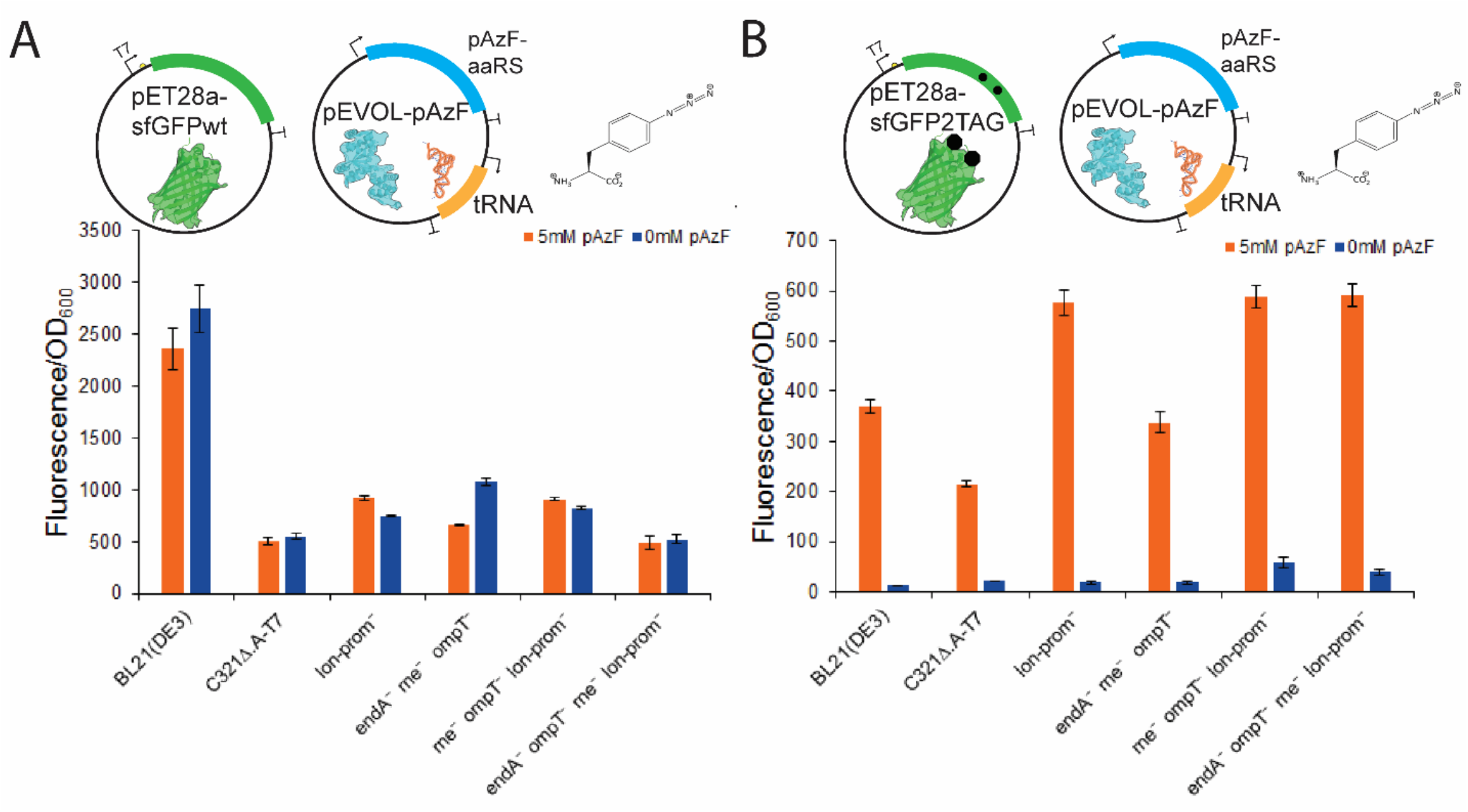
Expression of sfGFP in *C321.ΔA-T7* strains using a pET reporter plasmid. A) SfGFP-wt and the pAzF orthogonal translation system was expressed using a pET plasmid and pEVOL-pAzF, respectively. For all conditions 1 mM IPTG, 0.02% arabinose and 5 mM pAzF (orange bars) or 0 mM pAzF (blue bars) were added at OD_600_ 0.6-0.8. B) Expression of sfGFP with two amber codons at positions 190 and 212 in the presence (orange) or absence (blue) of 5 mM pAzF was tested with pEVOL-pAzF. For all panels error bars represent one standard deviation for biological triplicates and technical triplicates.

**Supplemental Figure 4.**
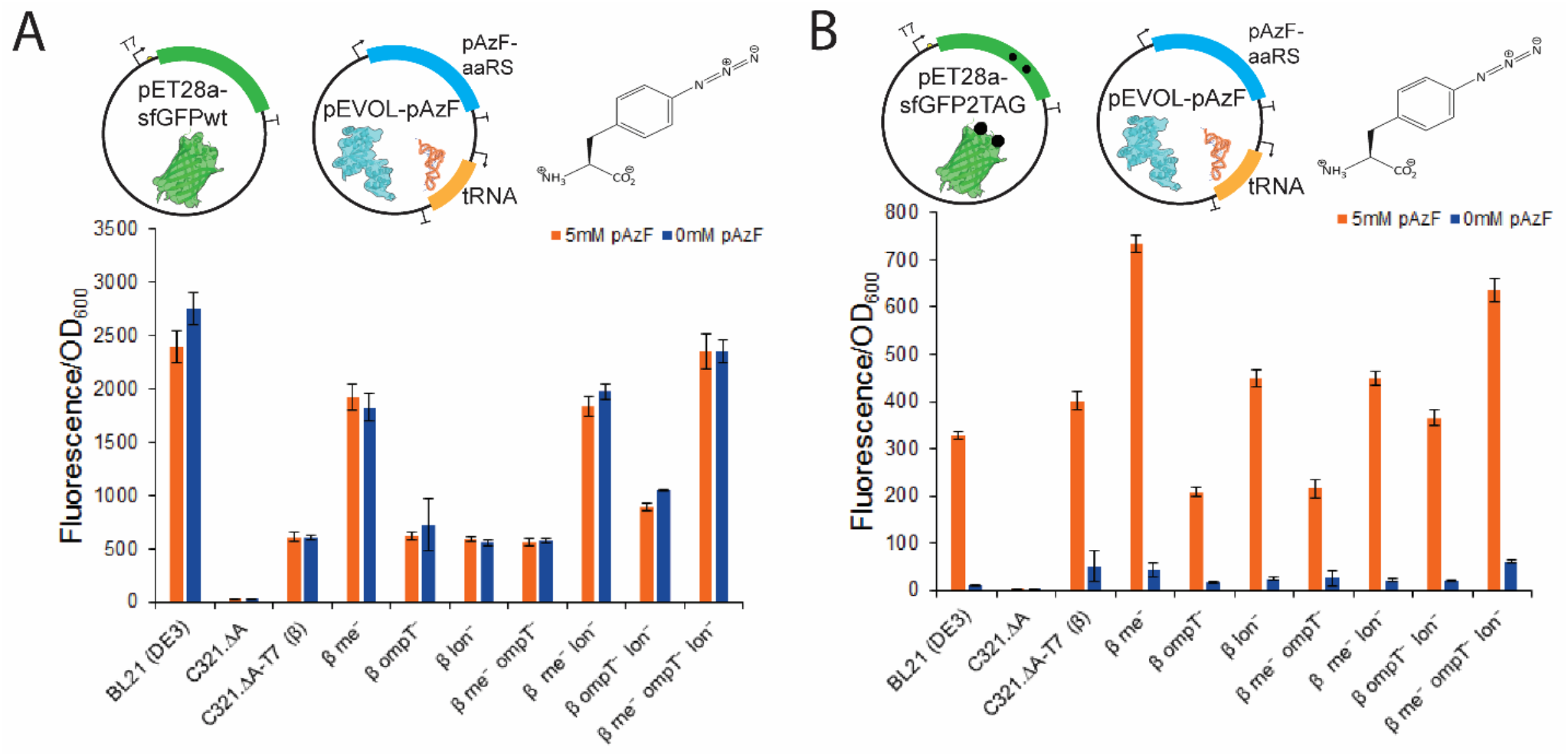
Expression of sfGFP in *C321.ΔA-T7* strains produced by first inserting T7RNAP, followed by the introduction of protein and nuclease mutations. A) Expression of sfGFP-wt was performed using pET28a-sfGFP-wt and pEVOLpAzF. For all conditions 1 mM IPTG, 0.02% arabinose and 5 mM pAzF (orange bars) or 0 mM pAzF (blue bars) were added at OD_600_ 0.6-0.8. B) Expression of sfGFP with two amber codons at positions 190 and 212 in the presence (orange) or absence (blue) of 5 mM pAzF was tested with pEVOL-pAzF. For all panels error bars represent one standard deviation for biological triplicates and technical triplicates.

**Supplemental Figure 5.**
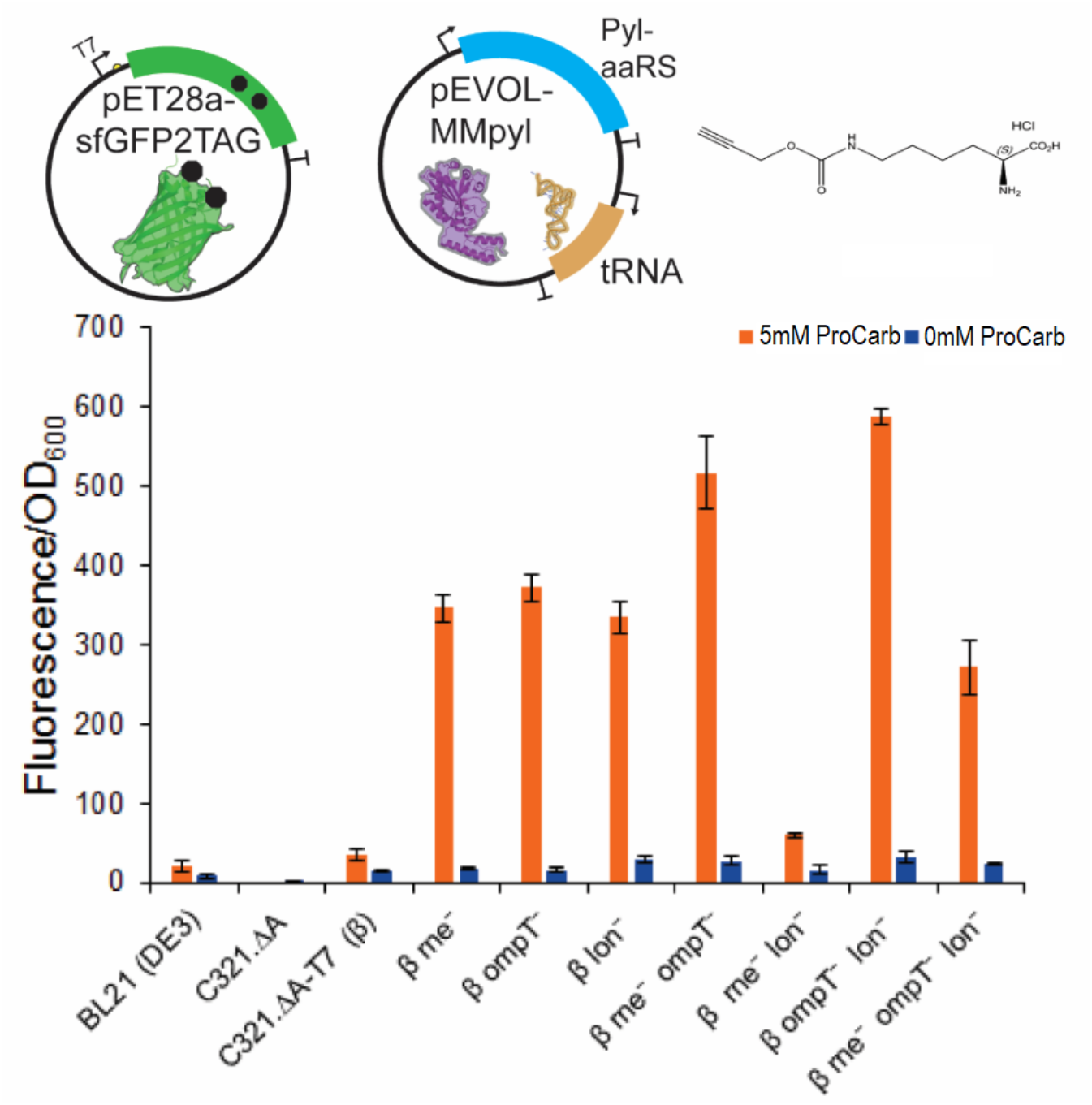
Expression of sfGFP-2TAG utilizing the pyrrolysine orthogonal translation system in β strains. Expression of sfGFP with two amber codons at positions 190 and 212 in the presence (orange) or absence (blue) of 5 mM ProCarb was tested with pEVOL-MMpyl. Error bars represent one standard deviation for biological triplicates and technical triplicates.

